# BNST GluN2D-containing NMDARs contribute to ethanol intake but not negative affective behaviors in female mice

**DOI:** 10.1101/2024.04.19.590258

**Authors:** Marie A. Doyle, Gregory J. Salimando, Megan E. Altemus, Justin K. Badt, Michelle N. Bedenbaugh, Alexander S. Vardy, Danielle N. Adank, Anika S. Park, Danny G. Winder

## Abstract

Alcohol use disorder (AUD) is a chronic, relapsing disease, highly comorbid with anxiety and depression. The bed nucleus of the stria terminalis (BNST), and *Crh*+ neurons in this region are thought to play a key role in chronic ethanol-induced increases in volitional ethanol intake. This role has been hypothesized to be driven by emergent BNST-dependent negative affective behaviors. Indeed, we report here that in female mice undergoing a home cage chronic drinking forced abstinence model (CDFA), excitatory transmission undergoes time-dependent upregulation in BNST *Crh*+ cells. Excitatory NMDA receptors (NMDARs) are a major target of ethanol, and chronic ethanol exposure has been shown to regulate NMDAR function and expression. GluN2D subunit-containing NMDARs have emerged as a target of interest due to their limited distribution and potential roles in affective behavior. We find that knockdown of dorsal BNST (dBNST) GluN2D expression significantly decreases ethanol intake in female, but not male, mice. While BNST *Grin2b* expression was significantly increased in protracted abstinence following CDFA, no differences in *Grin2d* expression were observed in dBNST or specifically in dBNST *Crh*+ neurons. Finally, to determine the impact of GluN2D expression on negative affective behaviors, open field, elevated zero maze, and forced swim tasks were used to measure anxiety- and depressive-like behaviors in constitutive and conditional BNST GluN2D knockout mice. Surprisingly, we find that deletion of GluN2D fails to alter negative affect in ethanol-naïve female mice. Together, these data suggest a role for BNST GluN2D-containing NMDARs in ethanol drinking behaviors but not abstinence from ethanol, highlighting potential sex differences and behavioral specificity in the context of AUD behaviors. Overall, these data further suggest roles for BNST synaptic signaling in volitional ethanol intake that are partially independent of actions on affective behavior.

## Introduction

Alcohol use disorder (AUD) is a chronic disease characterized by cyclic periods of alcohol intoxication, withdrawal, and anticipation (Koob and Le Moal, 1997). Though individuals with AUD may remain abstinent for years, relapse is common. One prominent theory proposes chronic ethanol-induced neuroadaptations within key brain regions as regulators of increased ethanol intake and seeking, hypothesized to be driven by negative affective behaviors that can emerge during subsequent abstinence (Zywiak et al., 1996, Dodge et al., 2005). Consistent with this idea, elevated levels of anxiety and depression are reported in patients with AUD (De Soto et al., 1985). However, traditional antidepressants are often ineffective in treating negative affect during withdrawal (Nunes and Levin, 2004), highlighting a need for novel therapeutics to better treat AUD.

Part of the extended amygdala, the bed nucleus of the stria terminalis (BNST) is a critical brain region for integrating negative affect and alcohol-related behaviors (Centanni et al., 2019a). This region is well known for its contribution to the regulation of stress- and anxiety-related behaviors, as chemogenetic activation of the BNST increases anxiety-like behaviors (Mazzone et al., 2018) while lesioning the BNST generally reduces them (Walker et al., 2009). Involvement of the BNST in affective behaviors translates onto abstinence from ethanol as well. In a mouse model of chronic drinking followed by forced abstinence (CDFA), capable of inducing anxiety- and depressive-like behaviors in protracted abstinence (Vranjkovic et al., 2018, Centanni et al., 2019b, Stevenson et al., 2009, Pang et al., 2013, Holleran et al., 2016), our lab has previously characterized behavioral and synaptic effects of withdrawal from continuous access to ethanol. Here, mice show increased BNST activity during prolonged abstinence, as measured by an increase in cFos-positive cells and glutamatergic transmission (Centanni et al., 2019b), together suggesting enhanced activity and glutamatergic input.

While previous work has characterized the BNST as a critical component of abstinence behaviors, the BNST has more recently been shown to orchestrate ethanol intake as well and is thought to play a key role in chronic ethanol-induced increases in volitional intake in AUD models. Altered BNST activity of specific neuronal cell types (Suresh Nair et al., 2022, Flanigan et al., 2023b, Companion and Thiele, 2018) and receptor function (Haun et al., 2020, Campbell et al., 2019) has been shown to modulate ethanol binge drinking. These data suggest a role for the BNST in not only abstinence phases of AUD but in intoxication phases as well, expanding the reach and influence of this region. However, molecular components of ethanol intake and their capacity to influence behavior across multiple phases, from intake to negative affect developed during abstinence from ethanol exposure, remain unclear.

*N*-methyl-D-aspartate receptors (NMDARs) are a major target of ethanol exposure (Lovinger et al., 1989). These heteromeric, ionotropic glutamate receptors are ethanol-sensitive and known to modulate the effects of ethanol in the BNST (Wills et al., 2012, Wills and Winder, 2013, Kash et al., 2009, Kash et al., 2008, Weitlauf et al., 2004). To date, research has primarily focused on GluN2A- or GluN2B-subunit containing NMDARs in mediating BNST ethanol sensitivity (Wills et al., 2012, Wills and Winder, 2013, Woodward et al., 2006, Weitlauf et al., 2005) due to their high prevalence relative to other subunits. The BNST also harbors GluN2D subunit-containing NMDARs (Salimando et al., 2020, Monyer et al., 1994, Sheng et al., 1994, Cull-Candy et al., 2001, Watanabe et al., 1993). GluN2D-containing NMDARs differ from other receptor compositions due to their unique biophysical properties, such as slower decay kinetics, magnesium insensitivity, heightened glutamate sensitivity, and low probability of opening (Traynelis et al., 2010). Their expression in the adult brain is highly restricted (Monyer et al., 1994, Sheng et al., 1994, Perszyk et al., 2016), highlighting GluN2D as a potential therapeutic target to consider in AUD treatment.

Our lab recently made significant inroads toward studying this subunit (Salimando et al., 2020), establishing the capacity for GluN2D expression to alter BNST synaptic transmission and negative affect. Specifically, BNST neurons expressing corticotropin-releasing factor (BNST^CRF^) of constitutive GluN2D knockout (KO) mice display increased excitatory postsynaptic currents (EPSCs) and decreased inhibitory postsynaptic currents (IPSCs), suggesting increased excitatory drive onto this neuronal subpopulation resulting from GluN2D manipulation (Salimando et al., 2020). Critically, constitutive GluN2D KO and BNST-specific conditional GluN2D knockdown male mice show enhanced anxiety- and depressive-like behaviors compared to same-sex littermate controls (Salimando et al., 2020, Yamamoto et al., 2017, Shelkar et al., 2019), defining BNST GluN2D-containing NMDAR mediated excitatory signaling as a potential component of negative affective behaviors. Despite this progress, the role and regulation of GluN2D-containing NMDARs in the context of ethanol intake and withdrawal as well as the behavioral consequences of BNST GluN2D deletion in female mice remain unstudied. Given the capacity for BNST GluN2D-containing NMDARs to alter glutamatergic transmission in the BNST and enhance negative affective behaviors in male mice, we hypothesized that GluN2D expression would alter both ethanol intake and negative affective behaviors and be regulated across abstinence periods. Thus, we used conditional deletion of GluN2D to assess the impact of BNST GluN2D expression on ethanol intake and negative affective behaviors in female mice and explored the regulation of *Grin2d* expression across behaviorally-relevant withdrawal time points during a mouse model of chronic alcohol consumption followed by forced abstinence (CDFA).

## Methods

### Animals

Male and female mice of at least 8 weeks of age were used in these studies. For gene regulation experiments, male and female C57Bl/6J mice (8 weeks of age) were purchased from Jackson Laboratory (#000664). Prior to the start of experiments, mice were allowed to habituate to the animal facility for at least 7 days. For GluN2D manipulations, conditional floxed GluN2D (FlxGluN2D) knockdown mice (*Grin2d*^tm1c[EUCOMM]Wtsi^, EMMA ID:04857) were re-derived and bred homozygously in house and constitutive GluN2D knockout mice were originally purchased from RIKEN Experimental Animal Division Repository (RBRC, #01840) and bred heterozygously in house. For identification of BNST corticotropin-releasing factor (CRF)-expressing cells in slice electrophysiology experiments, Crh-IRES-Cre and Ai14 (ROSA) mice were ordered from Jackson Laboratory (#012704 and #007914, respectively) and bred heterozygously. All transgenic lines were maintained on a C57Bl/6J background and genotyped at 3 weeks of age using the following primers (Salimando et al., 2020), respectively:

FlxGluN2D Forward: 5’- GTG TGA CCA GGA AGC CAC TT -3’
FlxGluN2D Reverse: 5’- TCC TTG ATC CCG TCC CTC AA -3’
GluN2D Primer 1: 5’- GCA GGC CCC TGC CTC CTC GCT C -3’
GluN2D Primer 2: 5’- CTG ACC TCA TCC TCA GAT GAG -3’
GluN2D Neo: 5’- TGG ATT GCA CGC AGG TTC TC -3’
Cre Forward: 5’- GGT CGA TGC AAC GAG TGA -3’
Cre Reverse: 5’- CCC TGA TCC TGG CAA TTT -3’
Ai14-ROSA 1: 5’- GGC ATT AAA GCA GCG TAT CC -3’
Ai14-ROSA 2: 5’- AAG GGA GCT GCA GTG GAG TA -3’
Ai14-ROSA 3: 5’- CTG TTC CTG TAC GGC ATG G -3’
Ai14-ROSA 4: 5’- CCG AAA ATC TGT GGG AAG TC -3’

All mice were group housed on a standard 12 hr light-dark cycle at 22-25°C with food and water available *ad libitum* and all behavioral assays took place during the light phase. All experiments were approved by the Vanderbilt University Institutional Animal Care and Use Committee (IACUC) and were carried out in accordance with the guidelines set in the Guide for the Care and Use of Laboratory Animals of the National Institutes of Health.

### Two-bottle choice tasks – Continuous access ethanol and chronic drinking followed by forced abstinence (CDFA)

To assess continuous access voluntary drinking, mice were singly housed with access to two 50 mL conical tubes fitted with sipper tops following established procedures (Centanni et al., 2019b, Vranjkovic et al., 2018, Holleran et al., 2016). Briefly, after habituating to the bottle setup for 7 days with only water in both bottles, ethanol-treated mice were provided one bottle containing an ethanol solution and the other containing water. Ethanol access began with an ethanol ramp: 3 days of 3% ethanol followed by 7 days of 7% ethanol. At the end of the ramp, mice had access to 10% ethanol for the duration of the ethanol access period, altogether totaling 6 weeks. Bottle and mouse weights were measured every 48-72 hours, and bottle placement was switched at the time of measurement to account for individual side bias. In studies assaying sucrose preference, at the end of the 6 weeks of ethanol access, the ethanol bottle was replaced with a bottle containing 1% sucrose for 7 days. Measurements were taken as described above.

In studies incorporating chronic ethanol drinking followed by forced abstinence (CDFA), ethanol access was described as above. At the end of the ethanol access period, the ethanol bottle was removed so that all groups had access to only water. Mice were then sacrificed during either acute (24 hours) or protracted (2 week) withdrawal, depending on the experimental setup.

### Whole-cell patch-clamp electrophysiology

Homozygous Crh-IRES-Cre mice were crossed with homozygous Ai14 mice to generate mice with tdTomato expression in corticotropin-releasing factor (CRF)-expressing cells. Female offspring were placed in the CDFA paradigm (described above) at 7-8 weeks of age. At either acute (24 hour) or protracted (2 week) abstinence time points, animals were transported to the laboratory and housed in a sound-attenuating cubicle for 1 hour prior to *ex vivo* brain slice preparation, as previously established (Salimando et al., 2020, Centanni et al., 2019b). Mice were deeply anesthetized with isoflurane and transcardially perfused with oxygenated, ice-cold N-Methyl-D-glucamine (NMDG)-based cutting solution (92mM NMDG, 2.5mM KCl, 20mM HEPES, 10mM MgSO_4_7H_2_O, 1.2mM NaH_2_PO_4_, 0.5mM CaCl_2_-2H_2_O, 25mM D-glucose, 3mM sodium pyruvate, 5mM sodium ascorbate, 2mM thiourea, 30mM NaHCO_3,_ and 5mM N-acetylcysteine) (Ting et al., 2018) before the brain was removed. The brain was hemisected, and 300 µm coronal sections were taken in an NMDG-based cutting solution using a VT1200S vibratome (Leica Microsystems). Slices containing the BNST were selected and transferred to a chamber containing warmed (34°C), oxygenated NMDG-based cutting solution for 10-15 minutes before being transferred to a holding chamber containing room temperature (25°C), oxygenated aCSF (124mM NaCl, 2.5mM KCl, 2.5mM CaCl_2_-2H_2_O, 5mM HEPES, 2mM Mg_2_SO_4_, 1.2mM NaH_2_PO_4_, 12.5mM D-glucose, 24mM NaHCO_3_, 0.4mM L-ascorbic acid) for at least 1 hour before the start of recordings.

After this recovery period, slices were transferred to a perfusion chamber and perfused with oxygenated, room temperature aCSF at a flow rate of 2mL/min for recording. Slices equilibrated for at least 10 minutes before the start of patching. BNST CRF neurons were identified by the expression of tdTomato. Glass pipettes (3-6 MΩ resistance) with silver chloride electrodes were filled with cesium gluconate internal solution (117mM Cs-gluconic acid, 20mM HEPS, 0.4mM EGTA, 5mM TEA, 2mM MgCl_2_, 4mM Na_2_ATP, and 0.3mM Na_2_GTP). Once access to a cell was achieved, the internal solution was allowed to diffuse for 10 minutes prior to recording. In voltage-clamp mode, spontaneous excitatory postsynaptic currents (sEPSCs) were recorded for minutes at -70mV, then the cell was slowly brought up to +10mV and allowed to equilibrate for minutes before spontaneous inhibitory postsynaptic currents (sIPSCs) were recorded for 4 minutes. Cells were excluded from the final analysis if access resistance changed by >20%. Signals were acquired with a Multiclamp 700B amplifier (Molecular Devices), digitized via a Digidata 1322A, and analyzed with pClamp 11.2 software (Molecular Devices) by measuring the peak amplitudes and frequencies of events over a 4 min period (in two, 2min bins).

### Viral-mediated gene transfer

#### Surgical procedure

Stereotaxic surgeries were performed following established procedures (Salimando et al., 2020). Mice were anesthetized with isoflurane (initial dose: 3%, maintenance dose: 1.5%) and received bilateral intra-dBNST infusions (0.3 μL) of rAAV5/CMV-Cre-recombinase (Cre)-GFP or rAAV5/TR-eGFP (University of North Carolina GTC Vector Core) at established coordinates (from bregma: A/P +0.14, M/L ±0.88, D/V –4.24, 15.03° tilt) (Salimando et al., 2020). Mice were allowed to recover for at least 21 days before the start of behavioral testing to allow for Cre-mediated gene deletion and the degradation of remaining GluN2D in transduced cells. Following the completion of behavioral assays, viral targeting was confirmed using standard histological methods (below).

#### Viral targeting validation

At the completion of behavioral studies, mice were sacrificed and perfused with 4% paraformaldehyde in phosphate buffered saline (PBS). Following cryoprotection in 30% sucrose-PBS, brains were sliced into 30 μm sections, and GFP labeling was used to confirm viral targeting. Mice with GFP expression outside of the BNST or with unilateral hits were excluded from analysis.

#### qPCR – BNST *Grin2d* knockdown validation

At least 3 weeks following stereotaxic surgeries for BNST GluN2D knockdown in male and female FlxGluN2D mice, mice were sacrificed, and the BNST was microdissected and stored at -80 °C until processing. RNA was isolated and purified using RNAeasy micro-columns (Qiagen), and cDNA was created using a QuantiTect Reverse Transcription Kit (Qiagen). Changes in *Grin2d* gene expression were determined via RT-PCR using SsoAdvanced Universal SYBR Green Supermix (BioRad) and PrimeTime qPCR Primers (IDT, *Grin2d*: Mm.PT.58.8048683, *Gapdh*: Mm.PT.39a.1, *Cre*: 487612563). Cre-recombinase (Cre) gene expression was used to confirm viral targeting of the BNST. All samples were run in triplicate and normalized to *Gapdh* before analysis using the ΔΔCt method (Doyle et al., 2020).

#### qPCR – BNST *Grin* gene regulation during abstinence

Female C57Bl/6J mice were individually housed in a two-bottle choice setup (CDFA, as described above) at the same time; however, ethanol access was staggered between acute and protracted withdrawal groups such that tissue could be collected from all groups on the same day at the end of the experiment. All groups were identically handled for the duration of the experiment. Following 6 weeks of two-bottle choice continuous ethanol access (10% ethanol) or water access (control condition), mice were sacrificed at either 24 hours or 2 weeks into a forced abstinence period. The BNST was microdissected and stored at -80°C until processing. RNA was isolated and purified using RNAeasy micro-columns (Qiagen), and cDNA was created using a QuantiTect Reverse Transcription Kit (Qiagen). Changes in *Grin2d* gene expression were determined via RT-PCR using SsoAdvanced Universal SYBR Green Supermix (BioRad) and PrimeTime qPCR Primers (IDT, *Grin1*: Mm.PT.58.33004583, *Grin2a*: Mm.PT.58.13771721, *Grin2b*: Mm.PT.58.42676841, *Grin2d*: Mm.PT.58.8048683, *Gapdh*: Mm.PT.39a.1). Cre-recombinase (Cre) expression was used to confirm viral targeting of the BNST. All samples were run in triplicate and normalized to *Gapdh* before analysis using the ΔΔCt method (Doyle et al., 2020).

#### RNAscope

RNAscope fluorescent *in situ* hybridization was performed as previously described (Salimando et al., 2020). Male and female C57Bl/6J mice underwent the CDFA protocol (described above), and tissue and blood were collected at the protracted (2 week) withdrawal time point. For RNAscope, the brains were removed, flash frozen in OCT compound, and stored at -80°C until sectioning. 16 µm coronal sections were collected using a CM3000 cryostat (Leica Microsystems), and sections containing the BNST were adhered to a warmed, charged glass microscope slice and immediately refrozen. Slides were stored at -80°C until tissue processing, following the Advanced Cell Diagnostics protocol for fresh frozen tissue (Multiplex Fluorescent Reagent Kit v1). Briefly, sections were first fixed with 4% paraformaldehyde and then dehydrated via a series of ethanol solutions. Next, experimental slices were incubated in the protease, probe (*Crh*: Mm-Crh-C2, *Grin2d*: Mm-Grin2d), and then amplification reagents, per standard kit instructions. Each slide contained 1 experimental slice and 1 control slice, which was treated as the experimental slice apart from receiving a negative control probe (*dapB*) instead of the *Crh* and *Grin2d* probes. Sections were also stained for DAPI before coverslipping. Next, the dorsal BNST of each slice was imaged as a z-stack using a 710 scanning confocal microscope (Carl Zeiss) at 20x magnification and images were analyzed using Fiji/ImageJ and Imaris. Negative control images were used to set brightness and contrast parameters for same-slide experimental images to control for autofluorescence. The dorsal BNST was analyzed as total *Grin2d* puncta counts, percent of BNST *Crh*-expressing cells co-expressing *Grin2d*, and average *Grin2d* puncta counts per BNST *Crh*-expressing cells.

#### Anxiety- and depressive-like behavioral testing

Measures of anxiety- and depressive-like behaviors were conducted within the Vanderbilt University Mouse Neurobehavioral Core. Mice were transferred to this facility and allowed to habituate for at least 7 days. Following habituation, animals were individually housed for at least 2 weeks before the start of testing. 5 days before this start, mice were handled by the experimenter to reduce experimenter-induced stress, as previously described (Olsen and Winder, 2010). All behavioral tests were conducted during the light cycle, and on test days mice were allowed to habituate to the behavior room for 1 hour prior to the start of experimentation. Tests were run in the following order: open field test, elevated zero maze, and forced swim test.

#### Open field test (OFT)

Mice were placed in ENV-S10S open field activity chambers (Med Associates) fitted with IR photograph-beam arrays for 1 hour. Locomotor activity was assessed by beam break counts, and anxiety measures were calculated as the percent of time spent in the center and periphery of the arena during the first 10 min of the session.

#### Elevated zero maze (EZM)

Mice were placed in the open quadrant of a custom EZM apparatus (34cm inner diameter, 46cm outer diameter, 40cm off the ground on four braced legs, and comprised of two open and two closed quadrants) for 5 minutes. Animal behavior was video recorded using an overhead camera and analyzed with ANY-maze software (Stoelting) as time spent in the open and closed quadrants.

#### Forced swim test (FST)

Mice were placed in a beaker containing room temperature (22-23°C) water for 6 minutes and video recorded. Time spent immobile (i.e. a lack of swimming or struggling movements) was hand-scored by experimenters blinded to experimental conditions.

### Statistics

Full statistical analyses and results are listed in Supplemental Table 1. All statistical analyses were performed using GraphPad Prism, and all values are represented as mean ± SEM. An unpaired t-test (two-tailed) was used to compare the means of two groups and a one-way analysis of variance (ANOVA) was used to compare the means of three groups, followed by a Tukey post-hoc test when appropriate. A two-way ANOVA was used when comparing two independent variables and a two-way ANOVA with repeating measures (RM) when two independent variables with repeated measures were compared, followed by a Sidak post-hoc test when appropriate, respectively. Males and females were combined for biochemical validation, but always analyzed separately for behavior. Significance was defined as *p<0.05 and **p<0.01.

## Results

### Glutamatergic input onto BNST^CRF^ neurons is increased in protracted, but not acute, withdrawal following ethanol availability

Our lab has previously demonstrated increased BNST activity and glutamatergic input during protracted abstinence from ethanol availability (Centanni et al., 2019b). However, the time course for the development of increased glutamatergic input remained unknown and potential changes in GABAergic input had not been previously assessed. To address this gap with a focus on BNST neurons expressing CRF (BNST^CRF^), previously implicated in both negative affect and stress-induced seeking behaviors (Silberman et al., 2013), CRF-Cre mice were crossed with an Ai14-ROSA reporter line to identify CRF-positive neurons with tdTomato expression. Female offspring were placed in the chronic drinking followed by forced abstinence (CDFA) paradigm. In this two-bottle choice task, mice were given continuous access to two bottles in their home cage, one containing water and the other ethanol. At the end of 6 weeks, the ethanol bottle was removed and mice were put into forced abstinence for either 24 hours or 2 weeks. Water controls had access to only water for the full duration of the experiment, and females were used due to the known sex difference in ethanol intake in this continuous access task and for consistency with previous work (Centanni et al., 2019b).

Importantly, there was no difference in ethanol intake between the 24 hour and 2 week withdrawal ethanol groups (F (1, 9) = 0.043, =0.84; Figure 1A). Animals were sacrificed following either acute (24 hour) or protracted (2 week) abstinence for *ex vivo* whole-cell patch-clamp electrophysiology. Spontaneous excitatory and inhibitory postsynaptic currents (sEPSCs and sIPSCs) were recorded from tdTomato-expressing CRF cells in the dorsal BNST (dBNST). Withdrawal groups were analyzed separately as tissue was collected at separate timepoints, with water control recordings taken at each timepoint and directly compared accordingly. In acute withdrawal, there were no differences observed in ethanol versus water mice in the frequency or amplitude of sEPSCs (frequency: t=0.44, df=24, p=0.66; amplitude: t=0.46, df=24, p=0.65; Figure 1B), suggesting no alteration of glutamatergic input at this time point. In protracted withdrawal, ethanol exposed mice displayed increased sEPSC frequency (t=2.38, df=22, p=0.027) but not amplitude (t=0.074, df=22, p=0.94; Figure 1C) compared to water controls, consistent with previous findings demonstrating increased glutamatergic input onto BNST^CRF^ neurons in protracted withdrawal (Centanni et al., 2019b) and the hypothesis of a hyperglutamatergic state in the BNST during abstinence. The frequency and amplitude of sIPSCs were not different at either acute (frequency: t=1.06, df=24, p=0.30; amplitude: t=0.014, df=24, p=0.99; Figure 1D) or protracted withdrawal (frequency: t=0.24, df=22, p=0.81; amplitude: t=1.09, df=22, p=0.29; Figure 1E) time points, suggesting no effect on phasic GABAergic transmission onto BNST^CRF^ neurons across the CDFA paradigm. These data replicate previous findings of increased glutamatergic input onto BNST^CRF^ neurons in protracted withdrawal from continuous ethanol access (Centanni et al., 2019b) and build on these existing data by characterizing increased BNST^CRF^ sEPSC frequency as only observed during protracted withdrawal and without concurrent changes in GABAergic transmission. Overall, these data describe the development of increased glutamatergic input across withdrawal, critical insight given the enhanced negative affect observed at this same time point. In previous studies we have provided evidence of the presence of synaptic GluN2D-containing NMDARs on the BNST^CRF^ neurons (Salimando et al., 2020). Therefore, given the observed regulation of glutamatergic transmission following ethanol access and the known ability of ethanol to modulate BNST NMDARs, we next sought to explore the potential role of GluN2D-containing NMDARs in ethanol-related behaviors.

**Figure 1:**
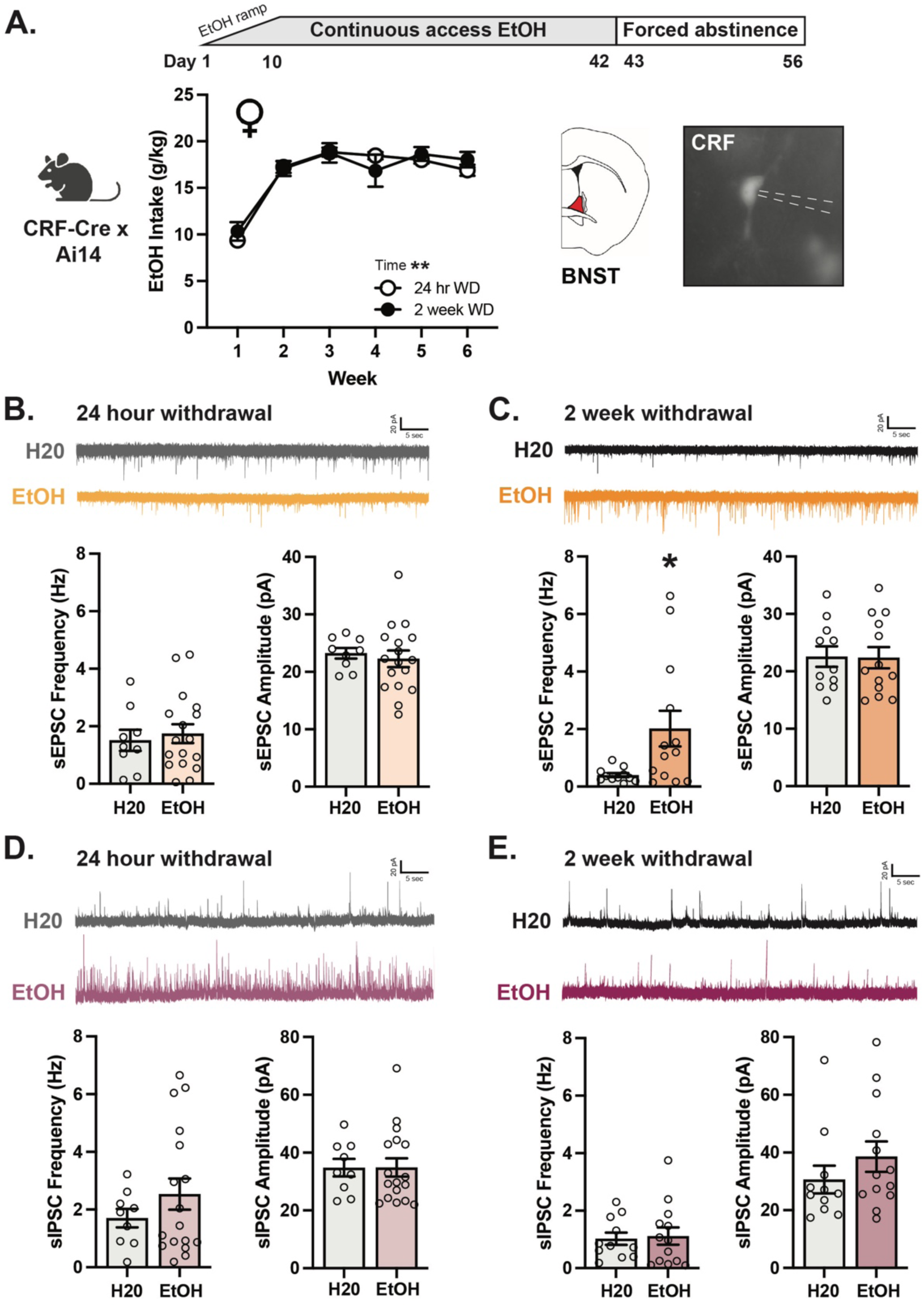
Increased glutamatergic transmission onto BNST^CRF^ neurons is increased in protracted, but not acute, withdrawal from continuous ethanol access. **A.** Timeline and schematic of recording from BNST^CRF^ neurons, with no differences in ethanol intake observed between withdrawal groups (n=5-6 mice, Two-way ANOVA with RM). **B.** Neither frequency nor amplitude of sEPSCs is altered in acute abstinence from ethanol compared to water controls (n=5-6 mice / 9-17 cells, unpaired t-test). **C.** Compared to water controls, there is a significant increase in sEPSC frequency during protracted abstinence in mice with previous access to ethanol, with no change in sEPSC amplitude (n=4-5 mice / 11-13 cells, unpaired t-test). **D.** The frequency and amplitude of sIPSCs are not altered in acute abstinence compared to mice with only access to water (n=5-6 mice / 9-17 cells, unpaired t-test). **E.** Protracted abstinence from ethanol access similarly does not alter sIPSC frequency or amplitude (n=4-5 mice / 11-13 cells, unpaired t-test). *p<0.05, **p<0.01. All data are represented as mean ± SEM.

### Conditional BNST knockdown of GluN2D decreased ethanol intake in a continuous access task

We first sought to characterize the impact of dBNST GluN2D-containing NMDARs on ethanol intake using a conditional GluN2D knockdown approach, as previous research has demonstrated the ability of the BNST to influence ethanol drinking behaviors. Specifically, modulation of BNST signaling (Campbell et al., 2019, Haun et al., 2020) and chemogenetic manipulation of BNST neuronal populations alter ethanol intake (Suresh Nair et al., 2022, Rinker et al., 2017, Companion and Thiele, 2018, Pleil et al., 2015, Flanigan et al., 2023b). Further, GluN2D knockout (KO) is known to alter BNST synaptic function (Salimando et al., 2020), suggesting the ability for expression of GluN2D-containign NMDARs to influence ethanol intake behaviors. To assess the ability for GluN2D deletion to alter ethanol intake, we again used a continuous access home cage task. To evaluate the impact of region-specific GluN2D expression, GFP- or Cre-encoding virus was bilaterally injected intra-BNST in male and female floxed GluN2D mice (Figure 2A). Viral-mediated knockdown of dorsal BNST *Grin2d* expression was validated by qPCR in a separate ethanol-naïve cohort, finding that Cre expression significantly decreased *Grin2d* expression compared to GFP controls (t=3.470, df=14, p<0.001; Figure 2B).

**Figure 2.**
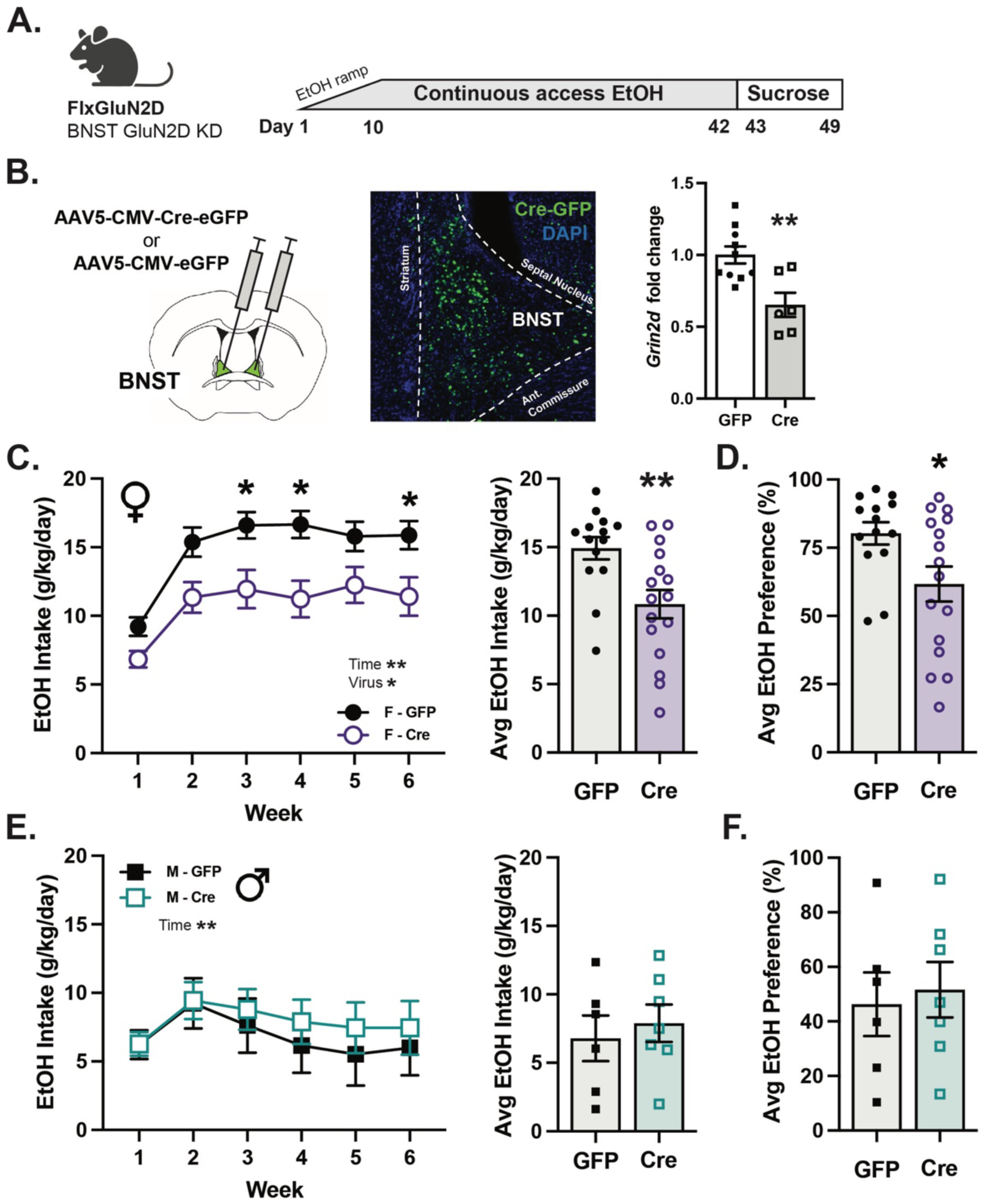
BNST knockdown of GluN2D significantly decreased ethanol intake in female mice. **A.** Experimental timeline. **B.** Cre recombinase significantly decreases *Grin2d* expression in the BNST of FlxGluN2D mice compared to GFP-expressing controls (females are represented as circles and males as squares, n=6-10, unpaired t-test). **C.** BNST knockdown of GluN2D significantly decreases ethanol intake in a continuous access task (left), with a significant difference in average ethanol intake (right) (n=13-16, Two-way ANOVA with RM with a Sidak post hoc test and unpaired t-test, respectively). **D.** Ethanol preference is also significantly decreased by BNST GluN2D knockdown in female mice (n=13-16, unpaired t-test). **E.** Ethanol intake is not altered by BNST GluN2D knockdown in male mice (n=6-7, Two-way ANOVA with RM and unpaired t-test, respectively). **F.** Male mice do not have altered ethanol preference as a result of BNST GluN2D knockdown (n=6-8, unpaired t-test).

During drinking experiments, males and females were analyzed separately due to pronounced sex differences in ethanol intake in this continuous access task, with females consuming significantly more ethanol than males (Centanni et al., 2019b). In female mice, there were main effects of time and virus on ethanol intake across weeks (time: F (5, 140) = 25.76, p<0.01; virus: F (1, 28) = 9.22, p<0.01) and a significant decrease in average ethanol intake (t=3.036, df=28, p<0.01) and preference (t=2.374, df=28, p=0.03) in BNST GluN2D knockdown mice compared to GFP controls (Figure 2C-D, Supplemental Figure 1). In male mice where ethanol intake was substantially lower, there were no differences observed between groups in ethanol intake behaviors (main effect of virus: F (1, 11) = 0.26, p=0.62; average intake: t=0.51, df=11, p=0.62) or in ethanol preference (t=0.28, df=11, p=0.79; Figure 2E-F and Supplemental Figure 1). There was no effect on overall fluid intake or on sucrose preference in either sex (Supplemental Figure 1). Together, these data suggest that ethanol intake can be regulated by the expression of BNST GluN2D-containing NMDARs in female mice.

### Withdrawal from continuous access ethanol does not alter BNST *Grin2d* expression in male or female mice

Given the increased glutamatergic input onto BNST^CRF^ neurons noted in protracted withdrawal (Figure 1C)(Centanni et al., 2019b) and a wealth of data demonstrating the effects of chronic ethanol exposure and withdrawal on NMDA receptor subunit expression, we next characterized BNST expression of key NMDA receptor subunit (*Grin)* genes at time points relevant to the development of negative affect phenotypes in CDFA. In the CDFA task, female C57Bl/6J mice were allowed to drink ethanol for 6 weeks, with controls only allowed access to water. No differences in ethanol intake were observed across ethanol withdrawal groups (F (1, 13) = 0.85, p=0.37; Supplemental Figure 2). Following 6 weeks of ethanol access, the ethanol bottle was removed from the cages so only water was available. Mice were sacrificed at either 24 hours (acute) or 2 weeks (protracted) withdrawal time points, and the BNST was collected via microdissection and processed for qPCR analysis. No differences were observed in NMDA receptor subunit *Grin1* (GluN1 subunit mRNA) expression (F (2, 23) = 2.17, p=0.14; Figure 3A), suggesting that overall expression of NMDA receptors may not be altered across withdrawal time points (Traynelis et al., 2010). BNST *Grin2a* (GluN2A subunit mRNA) was similarly unaltered across groups (F (2, 23) = 0.67, p=0.52; Figure 3A), consistent with previous *ex vivo* electrophysiology data indicating that this subunit is not a major driver of ethanol sensitivity in BNST (Kash et al., 2008, Wills et al., 2012) and suggesting the GluN2A subunit likely does not play a role in abstinence-regulated plasticity. There was a significant increase in BNST *Grin2b* (GluN2B subunit mRNA) expression (F (2, 23) = 5.10, p=0.015) when comparing the acute and protracted withdrawal time points (p=0.014), with a trend toward an increase when comparing water controls to the protracted withdrawal mice (p=0.054; Figure 3A). These data are consistent with BNST GluN2B-containing NMDARs showing a sensitivity to ethanol exposure (Kash et al., 2008, Wills et al., 2012). However, no significant differences were observed in *Grin2d* (GluN2D subunit mRNA) expression across any time point (F (2, 23) = 1.54, p=0.24; Figure 3A), indicating that *Grin2d* is likely not regulated in correlation with the development of ethanol-induced negative affective behaviors during CDFA.

**Figure 3.**
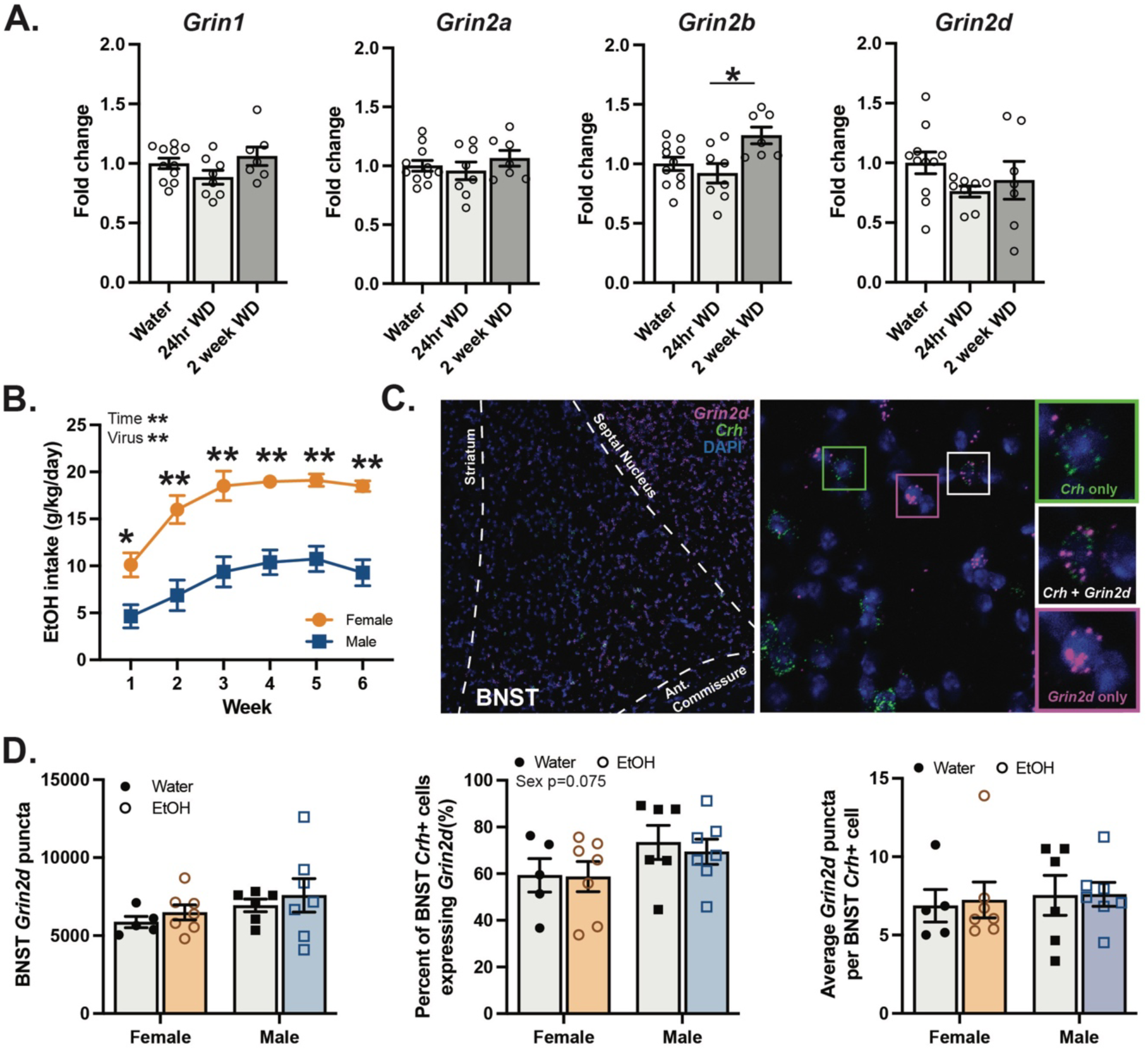
Neither BNST nor BNST^CRF^ expression of *Grin2d* is altered across withdrawal periods following exposure to continuous access ethanol. **A.** BNST expression of *Grin1*, *Grin2a*, and *Grin2d* are not altered across withdrawal periods compared to water controls; however, *Grin2b* was significantly increased in protracted withdrawal compared to acute withdrawal, with a trend toward an increase compared to water control mice as well (n=7-11, One-way ANOVA with a Tukey post hoc test). **B.** Female C57Bl/6J mice drank significantly more ethanol than male mice during the continuous access ethanol period (n=7, Two-way ANOVA with repeated measures with a Sidak post hoc test). **C.** Representative RNAscope image of the dorsal BNST. **D.** BNST and BNST^CRF^ expression of *Grin2d* is not altered in protracted abstinence in male or female mice, as measured as total BNST grin2d *puncta*, percent BNST^CRF^ cells expressing *Grin2d*, and average *Grin2d* puncta per BNST^CRF^ cells (n=5-7, Two-way ANOVA).

Though assessment of BNST *Grin2d* gene expression did not indicate regulation during withdrawal, qPCR may obscure changes occurring in specific subpopulations of neurons. A majority of BNST^CRF^ neurons express *Grin2d* (Salimando et al., 2020) and previous work identified effects of GluN2D deletion in BNST^CRF^ neurons but not randomly selected neurons (Salimando et al., 2020), indicating that BNST cell type specificity may be key to detecting relevant regulation of GluN2D-containing NMDARs. Further, BNST GluN2D knockdown increases anxiety- and depressive-like behaviors in ethanol-naïve male mice (Salimando et al., 2020), suggesting a contribution of GluN2D subunit expression to negative affective behaviors. To measure *Grin2d* expression specifically in the BNST^CRF^ subpopulation, fluorescent *in situ* hybridization (RNAscope) was used to determine the percentage of *Crh*-containing cells that expressed *Grin2d* as well as to quantify *Grin2d* puncta counts within *Crh*-containing cells. In this experiment, male and female C57Bl/6J mice underwent the CDFA paradigm, again with water controls with only access to water. All mice were sacrificed for tissue collection in protracted withdrawal (2 weeks). There was a main effect of sex on ethanol intake (F (1, 12) = 30.60, p<0.01), with females drinking significantly more ethanol than males (Figure 3B).

Following RNAscope analysis, no differences in dorsal BNST *Grin2d* puncta counts were detected across ethanol treatment (F (1, 21) = 0.82, p=0.38) or sex (F (1, 21) = 2.28, p=0.15; Figure 3D), consistent with no differences in BNST *Grin2d* transcripts assessed via qPCR analysis (Figure 3A). Next, analysis was focused on dorsal BNST cells expressing *Crh*. Here, no main effect of drug (F (1, 21) = 0.12, p=0.73) or interaction (F (1, 21) = 0.071, p=0.79) were observed in the percentage of *Crh*-containing cells that expressed *Grin2d*; however, there was a trend toward a main effect of sex (F (1, 21) = 3.52, p=0.075; Figure 3D). Further, no differences in *Grin2d* puncta counts within *Crh*-expressing neurons were detected across sex (F (1, 21) = 0.22, p=0.64) or ethanol treatment (F (1, 21) = 0.040, p=0.84; Figure 3D). Overall, these data indicate that *Grin2d* expression is not regulated in the BNST or in BNST^CRF^ neurons in protracted withdrawal from ethanol exposure during CDFA.

### BNST-specific GluN2D knockdown did not affect anxiety- and depression-like behaviors in female mice

Previous research has characterized negative affective behaviors in male constitutive knockout and conditional BNST knockdown mice, finding an increase in anxiety- and depression-like behaviors in these transgenic models (Salimando et al., 2020, Yamamoto et al., 2017, Shelkar et al., 2019); however, no studies have measured these behaviors in females. Therefore, we assessed the impact of GluN2D expression on negative affective behaviors in ethanol-naïve female mice using a behavioral battery including an open field task, elevated zero maze, and a forced swim test. To achieve BNST-specific gene knockdown, we again utilized a floxed GluN2D mouse line with a bilateral intra-BNST injection of Cre- or GFP-encoding virus to localize knockdown of GluN2D (Figure 4A). In ethanol-naïve females across all tasks observed, no differences in behavior were observed between Cre- and GFP-expression mice. Specifically, in the open field test, a main effect of time (F (11, 242) = 46.90, p<0.01) but not virus (F (1, 22) = 0.054, p=0.82) was observed (Figure 4B), with no difference in percent of time spent in the center of the field (t=0.53, df=22, p=0.60; Figure 4B). Similarly, the percent time spent in the open arm of an elevated zero maze was not different between Cre and GFP females (t=0.05, df=22, p=0.96; Figure 4C). Finally, time spent immobile was unchanged by BNST GluN2D knockdown in the forced swim test (t=0.51, df=22, p=0.61; Figure 4D). Overall, these data indicate that BNST GluN2D does not contribute to altered negative affect in female mice as measured in the assays conducted. As male constitutive GluN2D knockout (KO) mice had previously been found to display enhanced negative affect in these tasks (Salimando et al., 2020), we repeated this behavioral battery in female constitutive GluN2D KO mice to assess the effect of whole body GluN2D deletion on negative affect in females. There were significant main effects of genotype and time on locomotor activity (genotype: F (1, 18) = 8.797, p<0.01; time: F (11, 198) = 23.06, p<0.01) with decreased activity in GluN2D KO mice compared to wildtype littermates (Figure 4E), consistent with male data (Salimando et al., 2020). Time spent in the center of the arena was also significantly decreased in KO females (t=2.172, df=18, p=0.044, Figure 4E), suggesting increased anxiety-like behavior. However, GluN2D deletion did not alter time spent in the open quadrant of an elevated zero maze (t=0.2975, df=19, p=0.77) or time spent immobile in the forced swim task (t=1.582, df=19, p=0.13, Figure 4F-G), behaviors where an increase negative affective tendencies were detected in male GluN2D KO mice (Salimando et al., 2020). These data may suggest a sex difference in the contribution of BNST GluN2D expression to negative affective behaviors.

**Figure 4.**
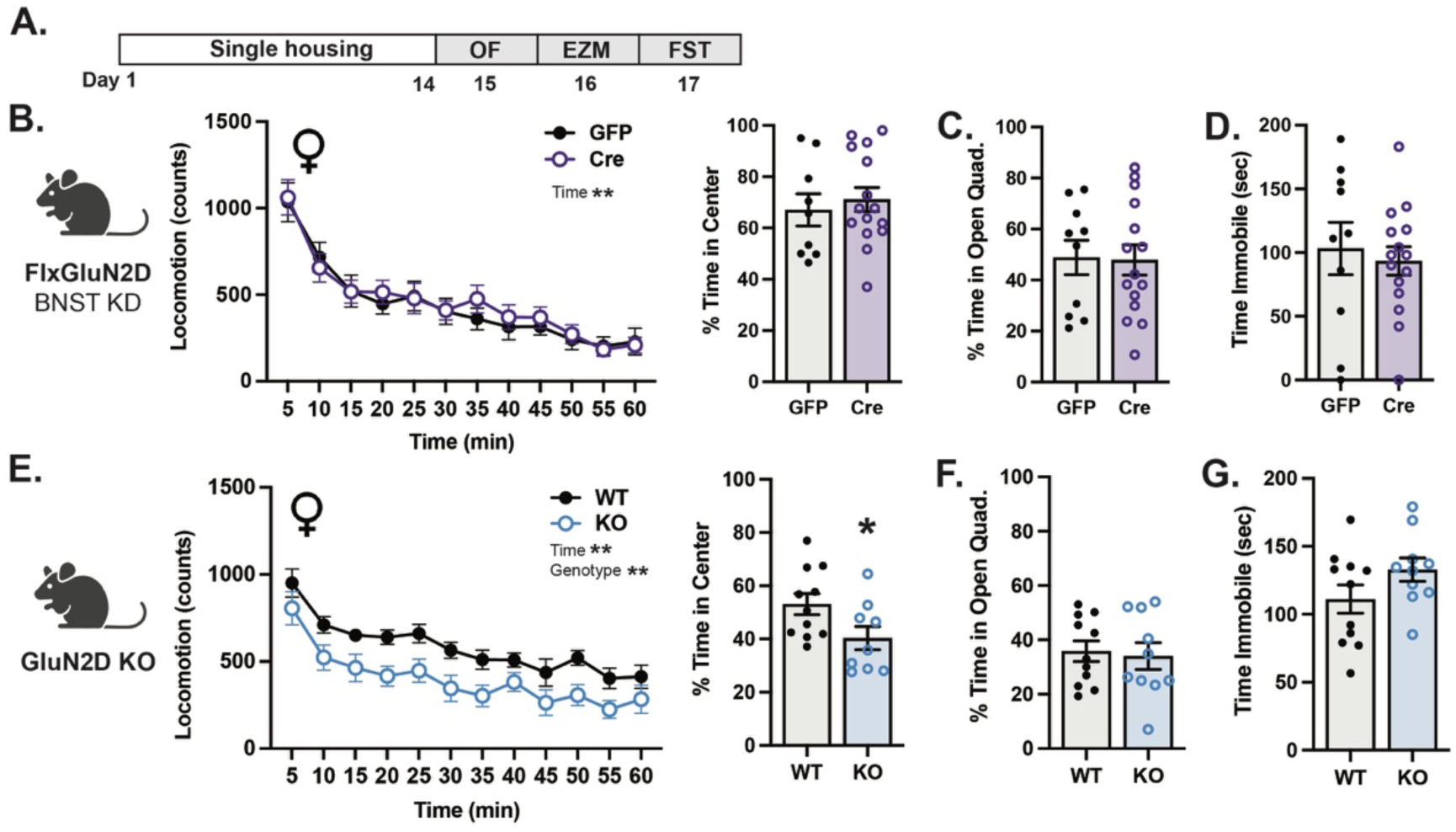
BNST knockdown of GluN2D does not alter negative affective behaviors in ethanol-naïve female mice. **A.** Experimental timeline. **B.** Knockdown of BNST GluN2D does not alter locomotor activity (left) or percent time spent in the center of the arena (right) (n=9-15, Two-way ANOVA with repeated measures and unpaired t-test, respectively). **C.** Percent time spent in the open quadrant of an elevated zero maze was not affected by knockdown of BNST GluN2D expression (n=9-15, unpaired t-test). **D.** Time spent immobile in the forced swim test is not affected by knockdown of GluN2D expression in the BNST (n=9-15, unpaired t-test). **E.** Constitutive GluN2D knockout deceases locomotor mice compared to wildtype littermate controls (left) and decreases percent time spent in the center of the arena (n=9-11, Two-way ANOVA with repeated measures and unpaired t-test, respectively). **F.** Percent time spent in the open quadrant of the elevated zero maze is not changed by GluN2D KO (n=9-11, unpaired t-test). **G.** Deletion of GluN2D does not alter time spent immobile in the forced swim test (n=9-11, unpaired t-test).

## Discussion

The bed nucleus of the stria terminalis (BNST) is a critical brain region for integrating alcohol-related and negative affective behaviors (Centanni et al., 2019a). Here, we demonstrate a role for BNST GluN2D-containing NMDARs in modulating ethanol intake in female mice. Specifically, knockdown of dorsal BNST (dBNST) GluN2D expression was capable of decreasing ethanol intake in female mice, without altering sucrose preference, and without an effect on male intake behaviors. To our knowledge, these studies are the first to demonstrate the influence of the BNST on moderate ethanol intake, link BNST NMDARs to volitional ethanol intake, and highlight a potential sex difference in the contribution of GluN2D-containing NMDARs in ethanol intake. However, though altered BNST^CRF^ neurons glutamatergic transmission was observed in protracted abstinence, BNST and BNST^CRF^ *Grin2d* expression was not altered across acute or protracted abstinence time points. Further, knockdown of dBNST GluN2D did not alter anxiety- and depression-like behaviors measured in ethanol-naïve female mice. Together, these data outline the contribution of BNST GluN2D-containing NMDARs in ethanol intake but likely not the development of negative affective behaviors during abstinence in female mice, defining potential sex differences and behavioral specificity in the context of AUD behaviors. Overall, these data further suggest roles for BNST synaptic signaling in volitional ethanol intake that are in part independent of actions on affective behavior.

### Contribution of BNST GluN2D expression to ethanol intake

Widely studied for its role in negative affect, the BNST also contributes to consummatory behaviors, including food (Wang et al., 2019, Williams et al., 2018, Jennings et al., 2013, Kocho-Schellenberg et al., 2014), saccharin (Kikuchi et al., 2023), and binge ethanol intake (Companion and Thiele, 2018, Suresh Nair et al., 2022, Flanigan et al., 2023b, Campbell et al., 2019, Haun et al., 2020, Flanigan et al., 2023a). In the present study, we found that decreased expression of GluN2D in the dBNST, a manipulation known to modulate BNST synaptic plasticity (Salimando et al., 2020), was sufficient to decrease ethanol intake in female mice during a continuous access task (Figure 2C). Ethanol is known to attenuate BNST NMDAR-dependent signaling (Wills et al., 2012, Weitlauf et al., 2004, Weitlauf et al., 2005, Wills et al., 2013, Kash et al., 2008). While the contribution of GluN2D subunit expression to these effects of ethanol in the BNST remains unstudied, experiments expressing recombinant NMDARs *in vitro* predict GluN2D-containing NMDARs to similarly be inhibited by ethanol (Smothers and Woodward, 2016). Our lab recently identified a deficit in BNST short-term potentiation of excitatory transmission recorded from constitutive GluN2D KO mice (Salimando et al., 2020), mirroring effects of acute ethanol application on BNST neurons (Weitlauf et al., 2004). Together, these findings are consistent with the role of BNST GluN2D-containing NMDARs in modulating ethanol intake.

### Sex differences in ethanol intake and the BNST

In this study, BNST-specific knockdown of GluN2D in adulthood decreased ethanol intake in female but not male mice (Figure 2C,E). A likely explanation for this distinction is the sex difference in ethanol consumed during the continuous ethanol access task (Figure 3B). Sex differences in ethanol intake in C57Bl/6J mice are observed across multiple home-cage intake paradigms (Centanni et al., 2019b, Levine et al., 2021, Middaugh et al., 1999, Hwa et al., 2011, Joffe et al., 2020), including this continuous access task. As male mice consumed less ethanol, subthreshold levels of intake may have failed to produce a behavioral effect in males. In future studies, intermittent ethanol access or a higher dose of ethanol may be necessary to drive sufficient levels of ethanol intake to engage the contribution of BNST GluN2D-containing NMDARs to ethanol intake behaviors in males. Additionally, estrogen is able to directly affect the expression of *Grin2d* (Watanabe et al., 1999). Therefore, neuroendocrine differences may drive differential gene expression or regulation of GluN2D in female mice compared to males. As estrous cycle was not tracked across drinking behaviors or at the times of tissue collection, future work is required to determine this impact on ethanol drinking behaviors.

### Altered BNST synaptic transmission during withdrawal in association with the development of negative affect

Our lab and others have previously characterized the development of negative affective behaviors during abstinence from continuous access ethanol intake in the CDFA model (Vranjkovic et al., 2018, Centanni et al., 2019b, Stevenson et al., 2009, Pang et al., 2013, Holleran et al., 2016). Acute withdrawal represents a key critical period in CDFA, as administration of ketamine at the onset of abstinence prevents the development of negative affective behaviors in protracted abstinence as well as increases the capacity for plasticity in the BNST in ethanol-treated mice (Vranjkovic et al., 2018).

In the present study we focused on BNST CRF-expressing neurons, as these cells have been linked to both negative affect and stress-induced seeking behaviors (Silberman et al., 2013) and are known to express *Grin2d* (Salimando et al., 2020). Using this approach we found that neither glutamatergic nor GABAergic transmission onto BNST^CRF^ neurons were altered in acute withdrawal (Figure 1B,D) but replicated enhanced glutamatergic transmission in protracted withdrawal (Figure 1C) (Centanni et al., 2019b). These data are consistent with increased cFos expression in BNST and BNST^CRF^ neurons during protracted abstinence (Centanni et al., 2019b). Together, these data define the timeline of altered glutamatergic transmission in withdrawal from ethanol and support the theory of a hyperglutamatergic state in the BNST during protracted abstinence, a time point relevant for the expression of negative affective behaviors.

### Regulation of *Grin* gene expression across periods of forced abstinence

GluN2D expression can be regulated by ethanol exposure, as *Grin2d* is upregulated in the hippocampus of patients with alcohol use disorder (Enoch et al., 2014) and alcohol-preferring rats express lower levels of amygdala *Grin2d* expression compared to saccharin-preferring rats (Augier et al., 2018). In the BNST, ventral BNST GluN1 and GluN2A protein expression were previously found to not be regulated in male mice during acute withdrawal from chronic intermittent ethanol vapor exposure, with GluN2B significantly upregulated compared to air controls (Kash et al., 2009), paralleling present mRNA findings in this continuous access paradigm in female mice (Figure 3A). However, we found that *Grin2d* expression was not altered across either withdrawal time point (Figure 3A). Because previously defined effects of GluN2D expression on BNST glutamatergic transmission were likely cell type-specific, as constitutive GluN2D KO mice show increased spontaneous and miniature EPSCs when recording from BNST^CRF^ neurons but not randomly sampled neurons (Salimando et al., 2020), we next used RNAscope to assay *Grin2d* transcripts specifically in BNST^CRF^ neurons. Though known sex differences in ethanol intake in C57Bl/6J mice were observed (Figure 3B-C), expression of *Grin2d* in BNST^CRF^ neurons was not altered across abstinence at time points critical to behavioral and electrophysiological effects or statistically different between sexes (Figure 3D). Together, these data suggest that GluN2D-containing NMDARs do not contribute to the development of negative affective behaviors during ethanol abstinence.

While a majority of BNST^CRF^ neurons express *Grin2d*, only a subset of *Grin2d*+ neurons are *Crh*+ (Salimando et al., 2020), allowing for the possibility of regulation within unexplored populations. Additionally, GluN protein levels or function were not assessed in the current study. Trafficking of GluN2D-containing NMDARs was not assessed in the present study and could account for GluN2D-driven changes in glutamatergic transmission without altered mRNA levels. For example, restraint stress can regulate synaptic vs. extrasynaptic localization of NMDARs (Tse et al., 2021) as well as increase membrane trafficking of NMDARs without altering protein levels (Yuen et al., 2009, Yuen et al., 2012). Phosphorylation of GluN2D subunits represents another potential regulatory step, as stress is known to modify GluN2B subunit phosphorylation (Ai et al., 2017).Therefore, regulation of *Grin2d* in other BNST neuronal subpopulations and protein-level regulation of GluN2D during abstinence periods cannot be ruled out.

### Sex differences in GluN2D regulation of anxiety- and depressive-like behaviors

Previous research has characterized the effects of constitutive GluN2D knockout and conditional BNST-specific knockdown as enhancing anxiety- and depressive-like behaviors in ethanol-naïve male mice (Salimando et al., 2020, Yamamoto et al., 2017, Shelkar et al., 2019). Expanding on previous work, we hypothesized that ethanol-naïve female mice would display similar effects of BNST GluN2D manipulation. In the present study, conditional BNST GluN2D knockdown and constitutive GluN2D KO in female mice did not show an effect of GluN2D expression on anxiety- or depression-like behaviors across a behavioral battery (Figure 4), failing to parallel findings from male mice with BNST-specific GluN2D knockdown (Salimando et al., 2020). These data may support a sex difference in the role of GluN2D expression in stress-and negative affect-related behaviors. Our lab has previously found BNST GluN2D to be downregulated following an acute restraint stressor in male, but not female, C57Bl/6J mice (Doyle et al., 2023). Further, male constitutive GluN2D knockout mice displayed decreased active coping bout behavior during repeated restraint stressors compared to same sex controls, an effect not observed in female mice (Doyle et al., 2023). Together, supported by the sexually dimorphic nature of the BNST as discussed above, these results may indicate a susceptibility to stress- and negative affect-behaviors conferred by GluN2D expression in male but not female mice.

## Conclusions

In summary, the present study supports a role for BNST GluN2D-containing NMDARs in the regulation of ethanol intake in a continuous access home-cage task but is likely not a driving force in the development of negative affect that arise during abstinence from ethanol exposure. These data additionally suggest a sex- and behavior-specific role for BNST GluN2D-containing NMDARs in AUD-related behaviors. Future studies will seek to more fully define the direct effects of ethanol exposure on GluN2D-containing NMDARs in the BNST and to better understand the potential role of GluN2D expression on sex differences in ethanol intake and negative affective behaviors.

## Supplemental figure legends

**Supplemental Figure 1.**
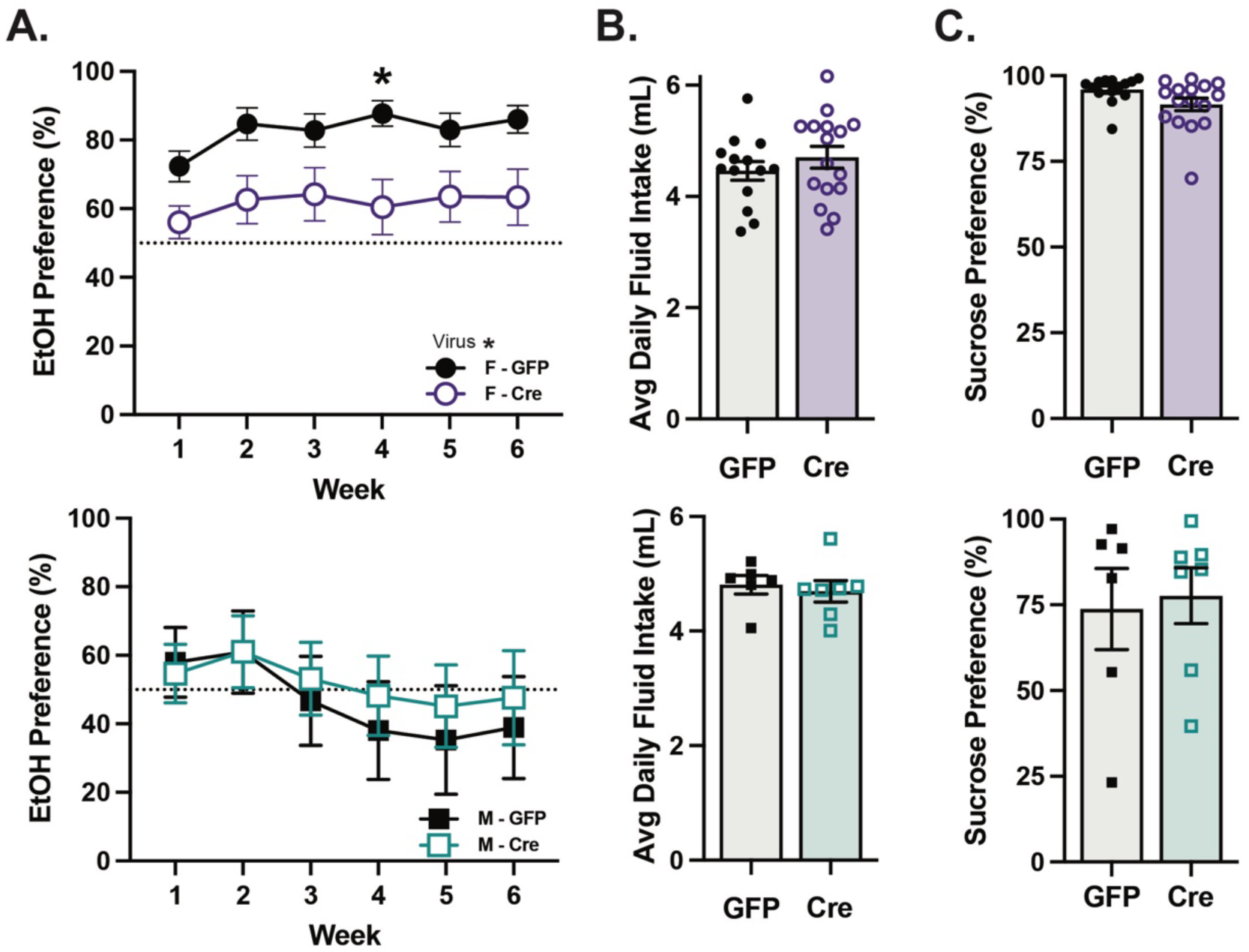
Ethanol preference but not daily fluid intake and sucrose preference are changed by BNST GluN2D knockdown in female mice. **A.** Ethanol preferences in male and female mice as a result of knockdown of BNST GluN2D expression (female n=13-16 and male n=6-7, Two-way ANOVA and unpaired t-test, respectively). **B.** BNST knockdown of GluN2D does not alter daily fluid intake in male or female mice (female n=13-16 and male n=6-7, unpaired t-test). **C.** Sucrose preference was not altered in either sex by BNST GluN2D knockdown (female n=13-16 and male n=6-7, unpaired t-test).

**Supplemental Figure 2.**
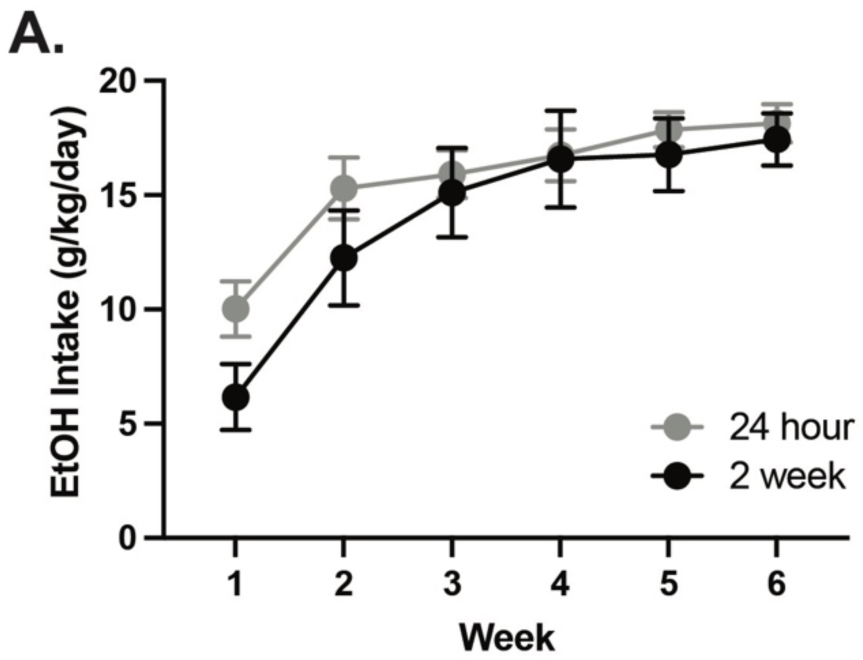
Ethanol intake was not different between withdrawal groups used for generating qPCR data. **A.** No difference in ethanol intake was observed between withdrawal time point groups (n=7-8, Two-way ANOVA with repeated measures).

## CRediT authorship contribution statement

**Marie Doyle:** Conceptualization, Methodology, Investigation, Formal analysis, Visualization, Writing - Original Draft, Project administration; **Gregory Salimando:** Conceptualization, Methodology, Investigation, Formal analysis, Writing - Review & Editing; **Megan Altemus:** Investigation, Resources; **Justin Badt**: Investigation, Validation; **Michelle Bedenbaugh:** Investigation, Resources, Writing - Review & Editing; **Alexander Vardy:** Investigation, Resources; **Danielle Adank:** Investigation, Writing - Review & Editing; **Anika Park:** Investigation, Validation; **Richard Simerly**: Funding acquisition, Resources; **Danny Winder:** Conceptualization, Writing - Review & Editing, Supervision, Funding acquisition, Project administration.

## Acknowledgements

Thank you to the Vanderbilt University Mouse Neurobehavioral Core for guidance in behavioral assays and to Bridget Morris and Laith Kayat for assistance with genotyping and colony maintenance.

